# Intraoperative contrast-enhanced ultrasound for cerebral glioma resection and the relationship with microvessel density

**DOI:** 10.1101/528281

**Authors:** Jia Wang, Xi Liu, Yilin Yang, Yunyou Duan

## Abstract

**Purpose:** We studied the value of intraoperative contrast-enhanced ultrasound (iCEUS) for real-time monitoring of resection of cerebral gliomas, and analyzed the relationship between CEUS parameters and microvessel density (MVD) of different pathologic grades of cerebral gliomas.

**Materials and Methods:** ICEUS was performed in 49 patients with cerebral gliomas. The enhancement characteristics of cerebral gliomas were observed before and after tumor resection. The number of microvessels was counted by immunostaining with anti-CD34. Differences in these quantitative parameters in cerebral gliomas were compared and subjected to a correlation analysis with MVD.

**Results:** The color Doppler flow classification within lesions were significantly different before and after iCEUS (p<0.05). The assessment of iCEUS parameters and tumor MVD showed that cerebral gliomas of different pathological grades had different characteristics. The time-to-peak (Tmax) was significantly shorter, the peak intensity (PI) and MVD were significantly higher in high-grade cerebral gliomas than in low-grade cerebral gliomas (p<0.05). According to the immunostaining, PI was positively (r=0.637) correlated with MVD and Tmax was negatively (r=–0.845) correlated with MVD.

**Conclusion:** ICEUS may determine the borders of lesions more clearly, indicate the microvascular perfusion in real time, and be helpful in understanding the cerebral gliomas grade.

## 1 Introduction

It has been confirmed that the tumor grade and complete tumor resection are essential for the prolonged survival of cerebral glioma patients[1]. Multiple studies have shown that intraoperative ultrasound (IOUS) is a valuable tool in tumor detection during cerebral surgery and it is becoming more widespread[2,3]. Although IOUS is excellent for tumor localization, little information is provided regarding microcirculation and perfusion dynamics in surgery[4]. According to our previous studies[5], the margin of the resection cavity is hyperechoic making the decision of further removal of tissue difficult if relied only on the IOUS. Recent advances in ultrasound technology, like 3D ultrasound and contrast-enhanced ultrasound (CEUS), have improved the image quality by increasing the sensitivity and alter the image contrast, especially CEUS, it is a dynamic and continuous way providing real-time view of vascularization and flow distribution patterns of different organs and tumors[6], but only few studies have been made to try to visualize brain lesions with CEUS[7], in fact, no guidelines exist on intraoperative CEUS (iCEUS) in neurosurgery.

In this study, we not only investigated the value of iCEUS in the real-time monitoring of cerebral glioma resection and judging whether the lesion was completely removed, but also we observed different morphologic and dynamic iCEUS patterns, analyzed the relationship between the quantitative parameters of iCEUS and tumor MVD in cerebral gliomas of different pathological grades, confirmed the reliability of iCEUS in assisting cerebral glioma grade.

## 2 Materials and methods

### 2.1 Research subjects

We performed intraoperative CEUS in 49 patients with cerebral gliomas (28 men and 21 women; age 17-68 years [mean, 42.3 years]) that were confirmed by preoperative magnetic resonance imaging (MRI) between October 2014 and December 2017 in the Department of Neurosurgery at Tangdu Hospital. The study protocol was approved by the Ethics Board of Department of Ultrasound Diagnostics, Second Affiliated Hospital of Air Force Medical University (Xin Si Road, No.569, Xi’an 710038, China). None of our patients had severe respiratory disease or heart disease. Informed consent was obtained from all participants in the study. All methods in this study were carried out in accordance with the approved guidelines.

### 2.2 Equipment and contrast agent

We used an SSD-α10 ultrasound system (Aloka, Tokyo, Japan) equipped with an intraoperative ultrasonic probe (UST-9120). The frequency was 3-6 MHz. As a ultrasound contrast agent (UCA), we used a second-generation UCA (SonoVue®, Bracco, Italy). Physiological (0.9%) saline (5 ml) and shaking were used to form a microbubble suspension of sulfur hexafluoride.

### 2.3 Operating procedure

All patients received general anesthetics and underwent microneurosurgical resection. A detailed medical history was obtained for each patient. The MRI enabled the location, shape, and borders of the glioma to be ascertained. The intraoperative probe was isolated with a sterile gum and gel and then was fixed on the operating table. After bone flap removal and the use of physiological saline as a coupling agent, we isolated the probe and placed it lightly on the dura or directly on the surface of the cortex. Two-dimensional (2D) ultrasound was applied to ascertain the extent and characteristics of the lesion and adjacent tissues and to measure the depth of the lesion from the cortical surface. Color Doppler was used to observe the blood supply of the lesion as well as the relationship between the lesion and important adjacent blood vessels. We scanned in multiple planes, such as coronal, sagittal and transverse, to determine the location of the lesion and the best approach for resection. Then, we selected the section with the largest lesion to do contrast-enhanced ultrasound. The mechanical index (MI) of SonoVue was 0.17–0.21, and the gain remained unchanged during the examination. The anesthesiologist injected SonoVue 2.4mL (5mg/mL) through the lower segment of the great saphenous vein and then injected physiological saline (10 ml) to flush the vein. The ultrasonic timer was then activated, and digital cine clips were stored continuously the tumor perfusion dynamics information in the US device. Then the lesion was removed by microneurosurgery, that is, using a microscope and cutting the cortex according to the gyri and sulci on the surface of the brain until the surgeon did a complete removal of the tumor. When the time of surgery, a first intraoperative qualitative analysis was performed, aimed at determining whether a contrast enhancement was detectable for every lesion and its characteristics of the contrast agent. After tumor resection, thorough hemostasis was performed. Hemostatic materials in the residual cavity were removed. The residual cavity was flushed with physiological saline. The probe was placed in the residual cavity, we observed the inner and outer portions of the residual cavity in multiple planes and determined the best view of the image. Then CEUS was performed again to observe the degree of enhancement of the residual cavity inner and outer portions. Data were also stored in the US device. Then, a second intraoperative qualitative analysis was performed, we selected the region that high or mild enhancement compared to brain parenchyma as regions of interest (ROI) and observed its contrast-enhanced features to determine whether a residual lesion was present. If contrast-enhanced features of ROI were similar to the lesion before resection, we thought this region was a suspicious residual lesion, the surgical team was asked to explore the region and repeat the resection. Residual tissue was sent for pathological examination. All data obtained by online and offline analysis were correlated with histopathology. Data were also stored in the US device for offline analysis.

### 2.4 Image analysis

We assessed the characteristics of the contrast agent using the time intensity curve (TIC) analysis provided by the corresponding software for the instrument. First, we examined the contrast imaging before tumor resection, selected the ROIs in the interior of the tumor (2-5 mm inside the tumor enhancement edge), and noted the tumor enhancement edge and normal brain tissue (10-20 mm outside the tumor enhancement edge). Each ROI was a 0.5-cm-diameter circle that was measured three times to obtain the average time-to-peak (T_max_) and peak intensity (PI) in the TIC analyses. Second two experienced sonographers graded the lesions according to the Adler classification of tumor blood flow as follows[7]: grade 0 (no blood-flow signals in the periphery or interior of the lesion); grade I (few blood-flow signals in the periphery and no or few (1–2) blood-flow signals in the interior); grade II (one main blood vessel with a length greater than the radius of the lesion or several small vessels extending from the periphery to the interior); and grade III (more than two vessels in the interior of the lesion or blood vessels communicating with each other that are interwoven into a network). Finally, we analyzed CEUS for the resected lesion and performed TIC analyses and calculated the T_max_ and PI to the suspicious residual lesion.

### 2.5 Determination of MVD

Postoperative biopsy specimens were fixed in 10% formalin, embedded in paraffin, and stained with hematoxylin and eosin. Immunohistochemical (IHC) analyses were carried out using anti-CD34 monoclonal antibody. We used the Weidner capillary counting method. Two experienced physicians performed a blinded analysis in which the field with the highest number of microvessels in a ×100 microscopic high-power field (hpf) was selected. Then, they randomly selected five microvessel-dense areas and counted the number of microvessels in a ×200 microscopic hpf. Individual or several brown-stained vascular endothelial cells and separate branches were classified as “vascular”. Vessels with a luminal diameter more than 8 times the diameter of red blood cells, necrotic areas, or sliced edges and a thick muscular layer of blood vessels were classified as “not vascular”.

### 2.6 Statistical analysis

SPSS v17.0 (SPSS, Chicago, IL, USA) was used for all analyses. The data are expressed as the means ± standard deviations. Independent sample *t*-tests were used to compare data between groups. The Spearman rank correlation test was used to assess correlations between parameters. Probability (p) values of < 0.05 was considered significant.

## 3 Results

### 3.1 Overall results

All patients underwent MRI to demonstrate that the lesions were single, had a depth of 0.9–5cm, and were located at different sites (frontal lobe, 17 cases; temporal lobe, 15 cases; parietal lobe, 10 cases; temporal and frontal lobe, 7 cases). According to the classification of cerebral gliomas (World Health Organization, 2007) [8] and postoperative pathological diagnosis, grade I, II, III, and IV cerebral gliomas were observed in 14, 12, 12, and 11 cases, respectively. Each patient was performed iCEUS twice before and after tumor resection, iCEUS yielded good results in patients through a bolus injection of contrast agent (2.4 ml), and no adverse events or side effects were observed. After iCEUS, we were able to directly visualize each of the 49 lesions before resection, the tumor border was clearer, the internal blood vessels of the tumor could be more clearly observed, and the relationships between the tumor and blood vessels, vascular-perfusion characteristics, and flow direction were better observed than before iCEUS (Fig 1). Before iCEUS, 46% of the color Doppler flow within the lesion was classified as grade III, whereas after iCEUS, 66% of the color Doppler flow within the lesion was classified as grade III (Fig 2).

**Fig 1.**
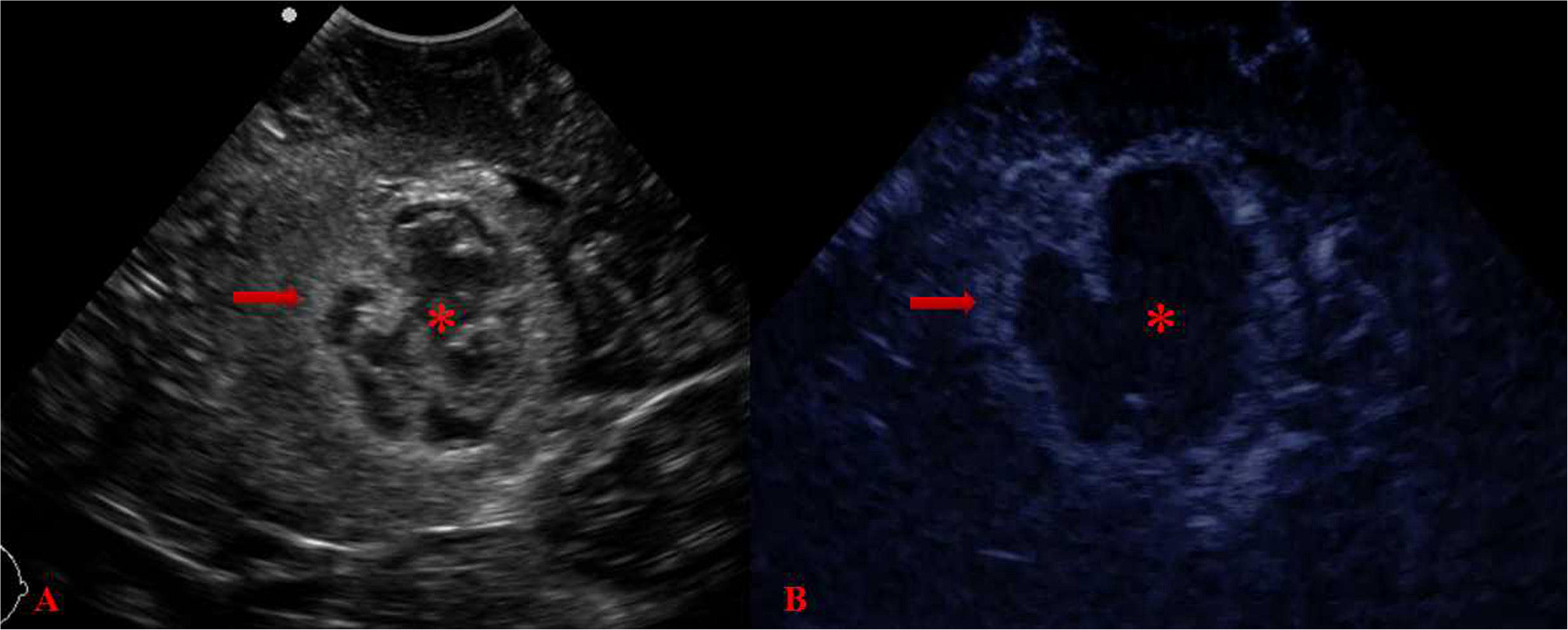
The lesion characteristics before and after CEUS. A) Before CEUS, the lesion was hyperechoic with irregular shape and indistinct margin (arrow), the internal echo was uneven (_*_), surrounding brain edema that was difficult to distinguish from the lesion; B) After CEUS, the lesion exhibited high enhancement (arrow), the border, internal echo (_*_) and boundary between surrounding brain edema and the lesion were more clear than before CEUS

**Fig 2.**
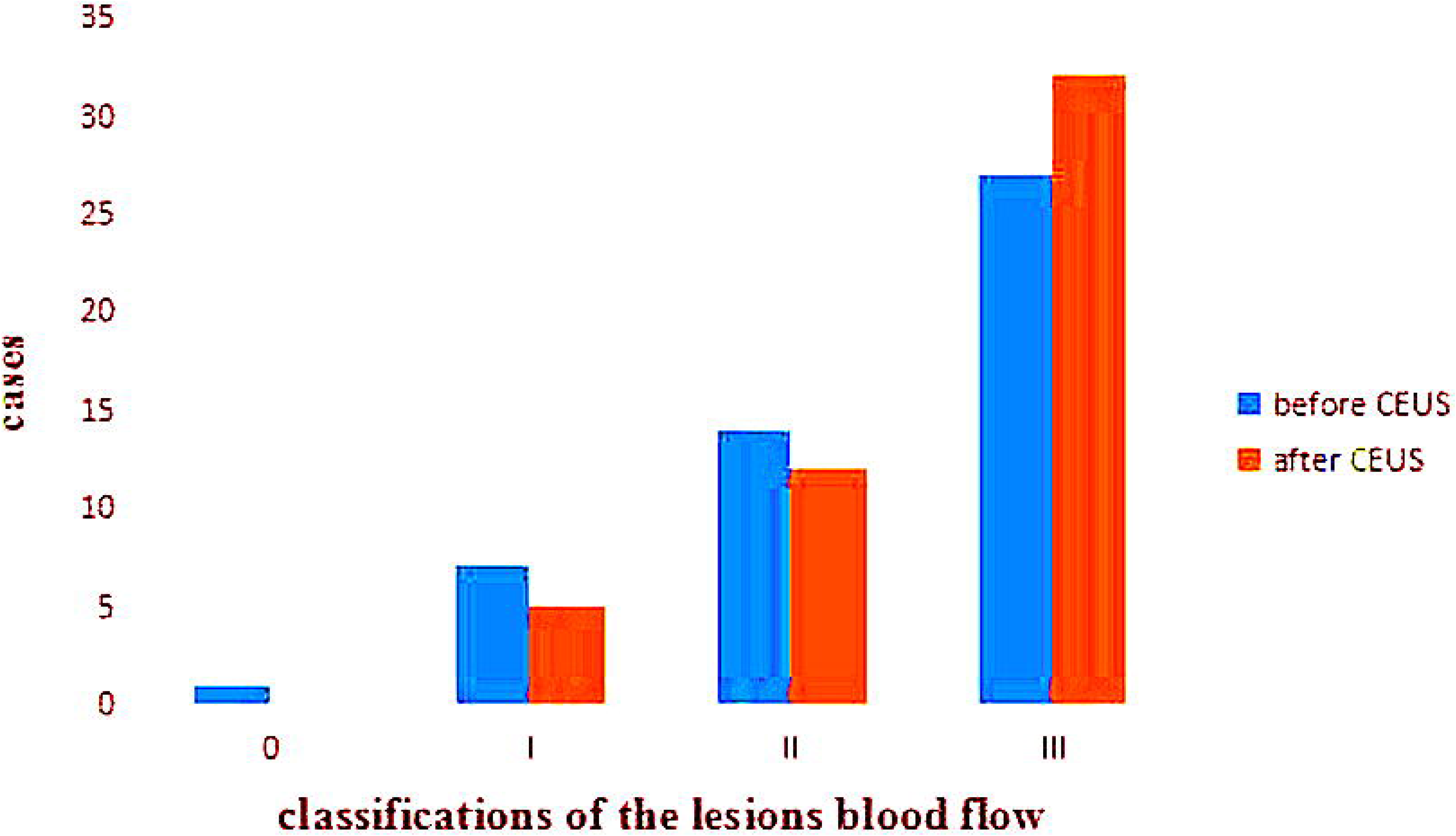
Profile of imaging classifications of the lesions blood flow before and after CEUS. Classifications of blood flow showed the lesions were classified as grade III after CEUS more than before CEUS

### 3.2 The iCEUS characteristics

Contrast-enhanced areas of lesions could be observed 16-27 s after the injection of contrast. Contrast-enhanced regions showed a rich blood supply, whereas non-contrast-enhanced regions showed a less rich blood supply, cystic degeneration, or necrosis. The iCEUS characteristic of normal brain tissue was “grid-like” with moderate enhancement. Peripheral edematous zones of lesions were observed at low contrast enhancement, and edema and tumor tissue could be distinguished. Different pathological grades of cerebral gliomas had different CEUS characteristics. Low-grade gliomas (LGGs) (grades I and II) exhibited a “lumpy” appearance and slight enhancement and could be differentiated from normal brain tissue (Fig 3). High-grade gliomas (HGGs) (grades III and IV) were also lumpy but exhibited significantly higher enhancement; additionally, the enhanced signal was uneven, thirteen lesions had no enhancement areas, all lesions had clear boundaries with surrounding tissue, (Fig 4).

**Fig 3.**
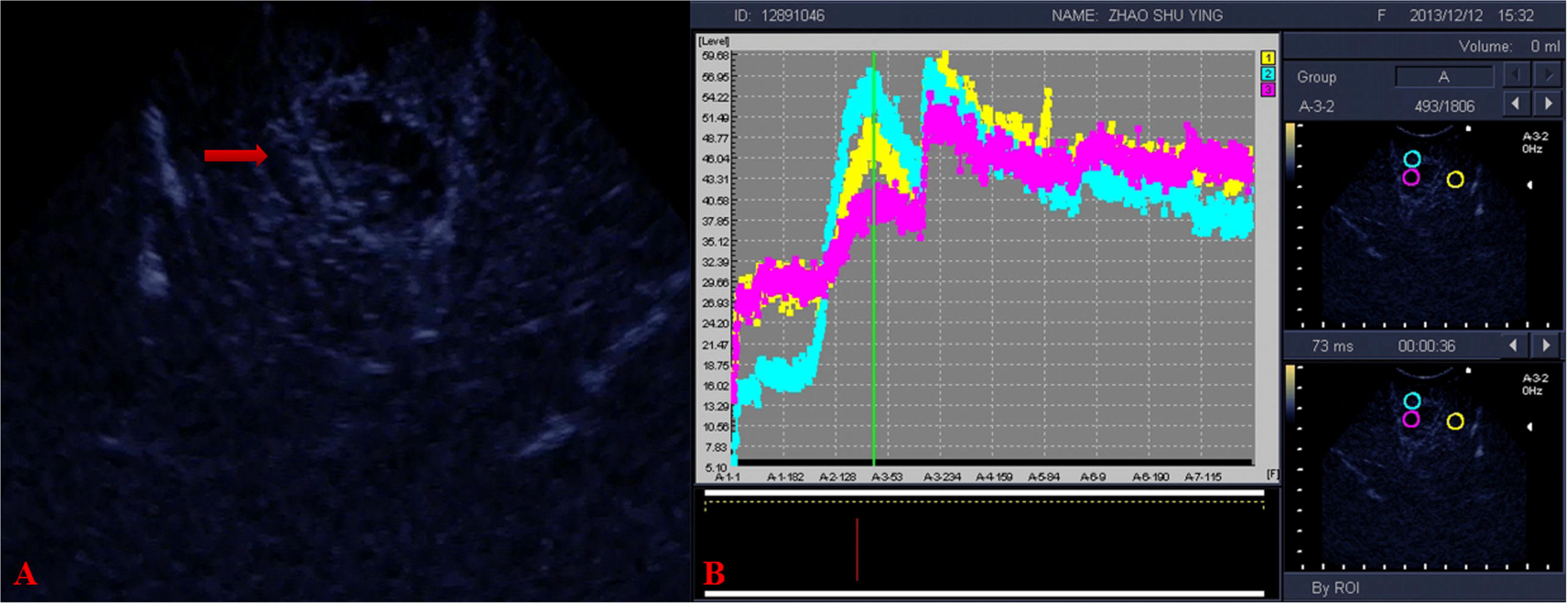
Contrast-enhanced ultrasound characteristics of a low-grade cerebral glioma. A) The lesion exhibited slight enhancement (arrow); B) Measures of T_max_ and PI of the ROI

**Fig 4.**
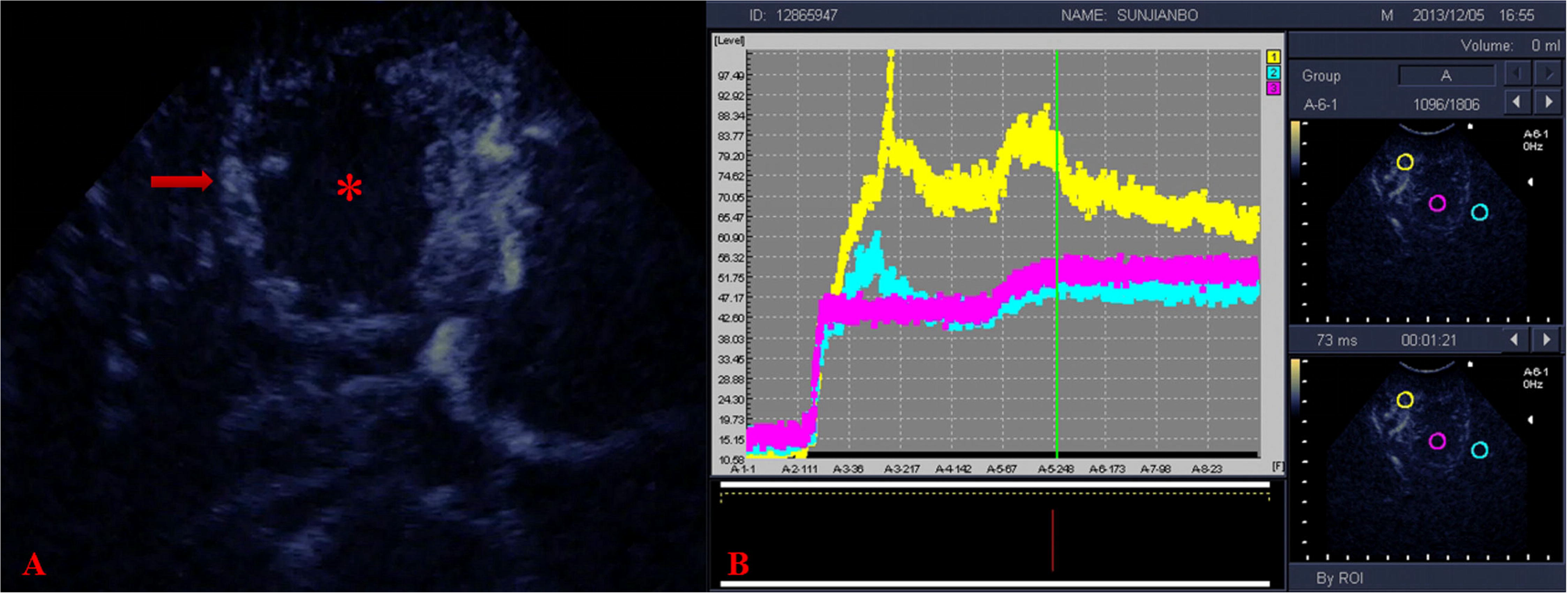
Contrast-enhanced ultrasound characteristics of a high-grade cerebral glioma. A) The lesion exhibited an uneven an uneven (_*_) and higher enhancement than surounding tissue (arrow); B) Measures of T_max_ and PI of the ROI

### 3.3 Histopathological diagnosis

The time intensity curve (TIC) analysis showed that the mean PI of normal brain tissue was 39.4±9.2dB, and the mean T_max_ was 48.8±8.6S. The mean PI of LGGs was 54.91±5.1dB, and the mean T_max_ was 42.7±5.4S. The mean PI of HGGs was 90.1±14.5dB, and the mean T_max_ was 30.6±7.2S. Comparison between LGG group and HGG group showed the mean PI of HGGs was significantly higher than that of LGGs, and the mean T_max_ was lower than that of LGGs; the differences between the two groups were significant (p_PI_=0.000 and p_Tmax_=0.000). Comparison between LGG group and normal brain tissue group showed the mean PI of LGGs was significantly higher than that of surrounding normal brain tissue (p_PI_=0.000), and the mean Tmax of LGGs was significantly lower than that of normal brain tissue (p_Tmax_=0.002). Comparison between HGG group and normal brain tissue group showed the mean PI of HGGs was significantly higher than that of surrounding normal brain tissue (p_PI_=0.000), and the mean Tmax of HGGs was significantly lower than that of normal brain tissue (p_Tmax_=0.000).(Table 1). Microvascular endothelial cells in cerebral glioma tissue stained positive with anti-CD34 antibody. The MVD was 36.1–78.9 (mean, 54.9±10.5) in 49 cases. The MVD increased as the degree of tumor malignancy. Grade I was observed in 14 cases, grade II in 12 cases, grade III in 12 cases, and grade IV in 11 cases, which corresponded to the mean MVD values of 43.8±4.5, 51.5±3.7, 59.6±7.3, and 67.8±4.9, respectively. The mean MVD of HGG was significantly higher than that of LGG, and the difference was significant (p<0.05) (Table 2). Analyses performed with the Spearman rank correlation test showed a positive correlation between PI and MVD (r=0.637, p=0.0001) and a negative correlation between T_max_ and MVD (r=–0.845 p=0.0001) (Fig 5 and Fig 6).

**Fig 5.**
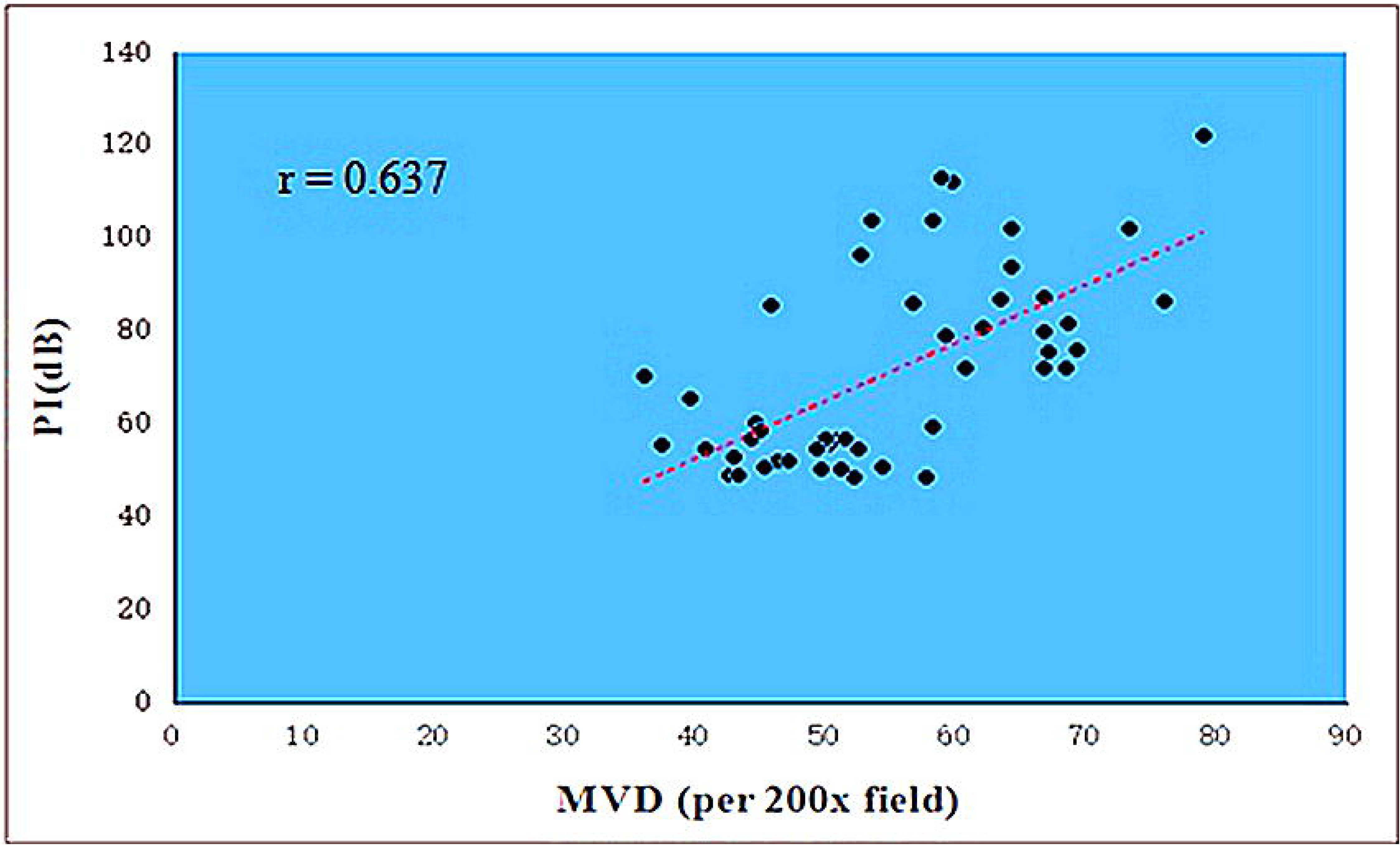
Correlation between PI and MVD (***x̄***±s) Showed a positive correlation between PI and MVD

**Fig 6.**
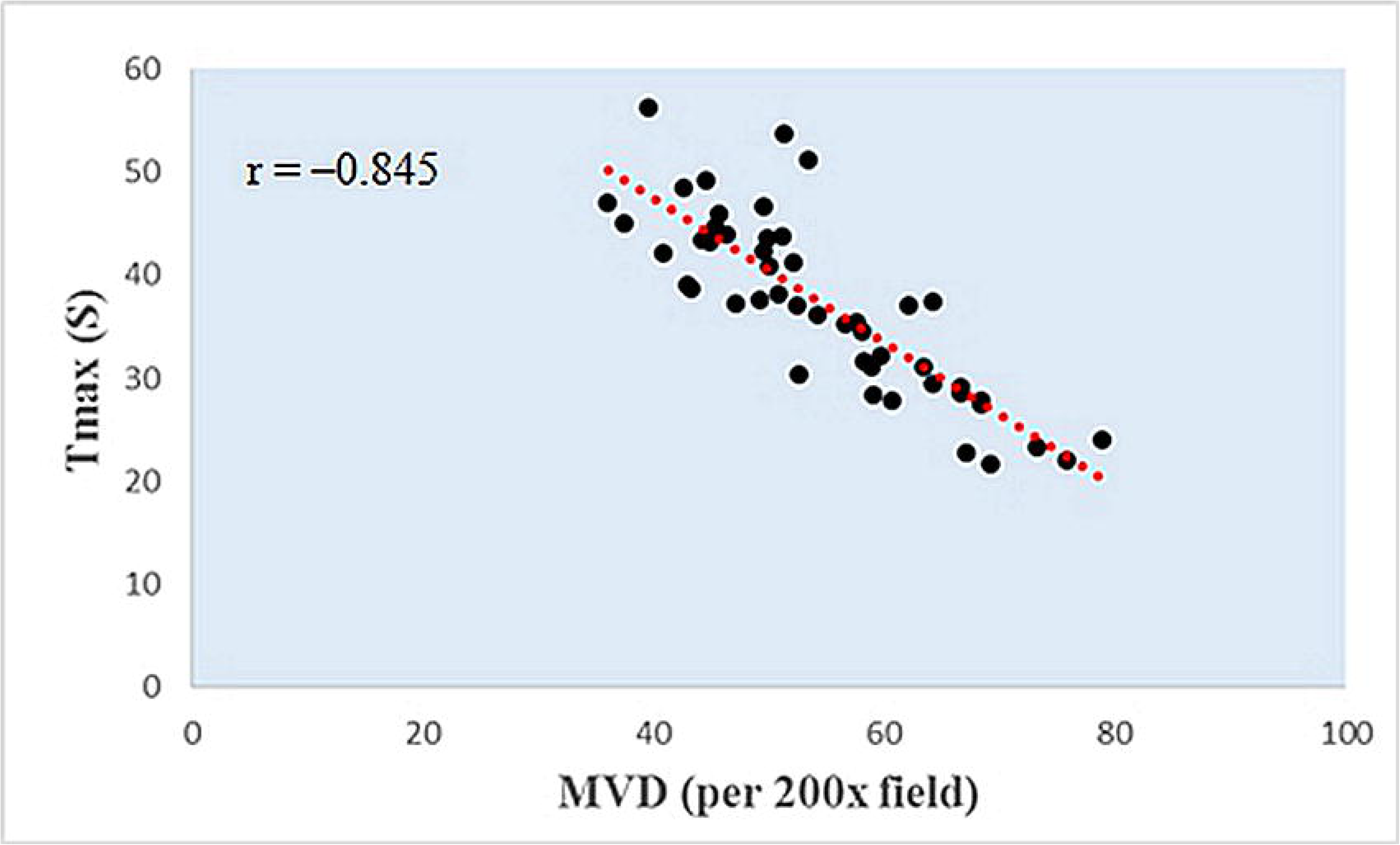
Correlation between Tmax and MVD (***x̄***±s) Showed a negative correlation between Tmax and MVD

### 3.4 Residual tumor tissue

Upon completion of iCEUS after tumor resection, nine residual cavities still had contrast-enhanced features that were observed preoperatively. Therefore, the surgical team performed further exploration and resection. In eight cases, the resected tissue was confirmed to be tumorous by pathological examination. In another case, the left edge of the residual cavity had a lumpy, high-contrast-enhanced appearance, and surgical exploration demonstrated the presence of a choroidal structure (Fig 7).

**Fig 7.**
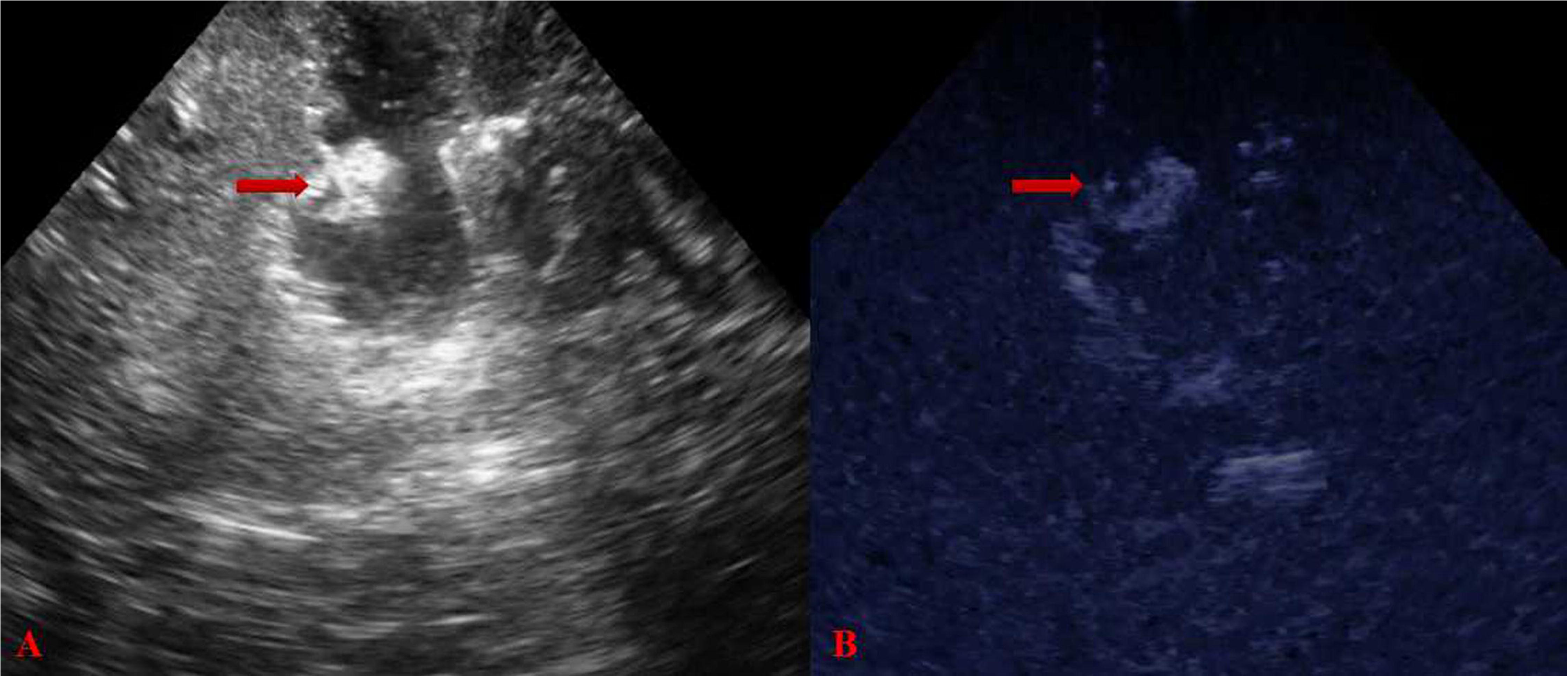
CEUS characteristics after tumor resection. A)The left edge of the residual cavity showed hyperechoic in IOUS (arrow); B) The left edge of the residual cavity with a “lumpy” high enhancement (arrow), which had been verified to be the choroidal structure by surgical exploration.

## 4 Discussion

The patient outcome of surgery, however, seems to be based on several factors, the pathological classification is important, especially for a variety of cerebral gliomas [9]. The degree of blood vessel growth is an important parameter in cerebral glioma classification, and tumor MVD is a reliable criterion for cerebral glioma angiogenesis. However, tumor MVD has a major limitation, namely, it cannot be used to detect tumor angiogenesis in vivo, and tumor specimens must be obtained [10]. Thus, identifying a noninvasive method to evaluate the MVD of tumors is important for improving free survival rates and ameliorating the quality of life in glioma patients.

IOUS is a readily repeatable, dynamic, and inexpensive procedure that can be performed at any time for a potentially unlimited number of times during surgery, but it provides little information regarding microcirculation and perfusion dynamics. Intraoperative MRI (iMRI) provides the surgeon with detailed image information; however, it requires a high degree of construction and logistics and is expensive [11]. A few studies have focused on intracerebral lesions with iCEUS or intraoperative 3D ultrasound to obtain information of tumor structures, differentiate between tumor and brain tissue, identify tumor remnants, provide the surgeon with information regarding dynamic changes that occur during surgery. Prada et al.[12,13] used iCEUS to intracerebral surgery showed that iCEUS is very useful in evaluating the location, defining the border, differentiating malignant and benign lesions and in maximizing tumor resection, thus improving free survival rates in cerebral glioma patients. Wang et al [14] evaluated the feasibility and value of iCEUS in different pathological grades of cerebral glioma apparently without obtaining iCEUS characterization for these lesions. Unsgaard et al [15,16] reported that intraoperative 3D ultrasound have a significantly better agreement with histopathology than navigated MRT1 for low-grade astrocytomas. However, few studies have been conducted different grades of malignancy in cerebral glioma surgery. Our findings showed that iCEUS increased the imaging sensitivity of the blood supply and microvessels of cerebral gliomas. It could be used to observe perfusion in the microcirculation of cerebral gliomas, thereby enabling its quantification to reflect blood-vessel growth in the cerebral glioma, which might relate to pathological grades of cerebral gliomas.

In general, the greater the degree of malignancy, the more rapid the growth of the tumor [17]. This scenario leads to the tumor needing a larger blood supply, which results in a greater number of blood vessels per unit volume. This situation leads to more obvious enhancement of contrast, and the degree of enhancement can reflect tumor MVD. Our study showed that the iCEUS characteristics of normal brain tissue and tumor tissue were markedly different. However, the iCEUS characteristics of normal brain tissue and those of edematous brain tissue were similar, showing grid-like and moderate enhancement. Cerebral gliomas apply pressure to surrounding brain tissue, and damage to the blood-brain barrier causes an increase in capillary permeability, with water leakage leading to cerebral edema; however, the number and distribution of microvessels are not significantly different [18]. ICEUS of HGGs appeared lumpy and showed significantly higher enhancement than surounding tissue. This feature illustrated the high degree of microbubble accumulation in the tumor, suggesting that the tumor blood supply was rich, and the microvascular distribution was clearly different from that in normal brain tissue. Thus, iCEUS indirectly reflected the high degree of proliferation of tumor blood vessels. ICEUS of LGGs also appeared lumpy but enhancement was slight, suggesting that the degree of proliferation of tumor blood vessels was lower than that observed in HGGs. If the internal blood supply to a tumor is short or interrupted, the local tumor tissue can undergo liquefaction necrosis. In the present study, 13 HGG cases showed homogeneous high echo by IOUS, but showed no enhancement areas by iCEUS, suggesting that these non-enhanced areas may have been necrotic. Surgery to remove this tissue showed jelly-like, amorphous material, and postoperative pathology confirmed liquefaction necrosis. After tumor removal we performed iCEUS in order to highlight tumor remnants, thus possibly maximizing resection. In 9 cases we visualized contrast enhancement areas, thus, the surgical team was asked to explore the suspicious lesions. Except for one case with a confirmed choroidal structure, eight cases were confirmed as residual tumors by pathological assessment. Consequently, once enhanced, the tumor is highlighted and reveals other specific characteristics. These findings might possibly be related to their grade.

In this study, we were able to directly visualize each of the 49 lesions before resection with iCEUS, and found that the different pathological grades of cerebral gliomas had different patterns of iCEUS enhancement. ICEUS-TIC analyses of cerebral gliomas reflected changes in flow velocity and flow volume of microbubbles in tumor vessels with time, and showed a very good correlation with histopathology[14]. This confirms once more the reliability of iCEUS in assisting tumor grading. Thus, CEUS, Tmax, and PI can indirectly be used to reflect the MVD of cerebral gliomas. We showed that T_max_ was shortened and PI was increased with a reduction in the degree of tumor differentiation and that the differences between the LGG and HGG groups were significant. IHC staining and quantitative analyses showed that the MVD in tumor tissue increased gradually with a reduction in the degree of differentiation of cerebral gliomas and that there were significant differences between the groups. Thus, the Tmax, PI, and MVD of cerebral gliomas may be connected in some manner. We found that T_max_ was negatively correlated with MVD. Thus, a lower degree of differentiation of cerebral gliomas was associated with a greater number of blood vessels per unit volume of the tumor. Therefore, the greater the amount of contrast agent that enters the tumor per unit volume of the tumor in in a given unit of time, the more significant the enhancement and, as a result, the easier it is to reach the peak value with a shorter T_max_.

Compared with the results from other studies [13, 19], our results showed that the start time of enhancement and T_max_ of cerebral gliomas were longer. This disparity may be associated with the path of injection of the contrast agent. We used a bolus injection of contrast agent through the lower segment of the great saphenous vein, but a more common path is through an elbow vein. Moreover, we observed iCEUS characteristics in intracranial tissue, and therefore the length of the circulation path of the contrast agent was increased, and the imaging time was prolonged. Furthermore, one major limitation of this study is related to the iCEUS technique. It is mandatory to accurately scan the lesion with IOUS before performing iCEUS to evaluate the more significant portion of the lesion and to obtain as much information as possible with regard to the timing, such as the tumor depth and appropriate imaging plane. Finally, another limitation is related to the experience of the examiner, and UCA injection has to be strictly synchronized with the start of the timer to achieve an accurate calculation of UCA transit time. To obviate such drawbacks, it is advisable that the examiners who perform iCEUS be trained, required close cooperation between neurosurgeons and ultrasound physicians, and both must be experienced in localizing anatomical structures and with the operative protocol. Of course iCEUS cannot be considered as an alternative to histological examination, which remains the gold standard for diagnosis.

With the development of ultrasonic technology, we believe that iCEUS is definitely a methodology to further understand and develop in cerebral glioma surgery and the expected results will leading to better treatment for the patients.

## 5 Conclusions

Performing iCEUS during cerebral glioma removal could provid dynamic and continuous real-time imaging and quantitative data analysis of different pathological grades of cerebral gliomas, and close to the completion of the resection. In addition, the quantitiative CEUS parameters were closely related to the MVD in different pathological grades of cerebral gliomas, might corroborate histological diagnosis, be helpful during surgery in differentiating malignant and benign cerebral glioma and refining surgical strategy.

## Acknowledgments

The authors gratefully acknowledge all the radiologists who helped with this survey and with the preparation of the manuscript. This manuscript was proofread by a native English-speaking professional with a science background at NPG Language Editing.

