## Supplementary material for "Intraoperative contrast-enhanced ultrasound for cerebral glioma resection and the relationship with microvessel density": table

**Table 1 Comparison of quantitative parameters of contrast-enhanced ultrasound in normal brain tissue, low-grade gliomas (LGGs), and high-grade gliomas (HGGs) (**
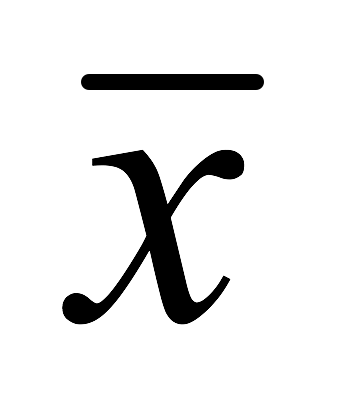
**±s)**

| **Tissue** | **Cases** | **PI (dB)** | **T_max_ (S)** |
| --- | --- | --- | --- |
| Normal brain tissue | 49 | 39.4±9.2#Δ | 48.8±8.6#Δ |
| LGGs | 26 | 54.9±5.1* | 42.7±5.4* |
| HGGs | 23 | 90.1±14.5 | 30.6±7.2 |

PI, peak index; T_max_, time-to-peak.

Comparison between groups: LGG and HGG *p<0.05; normal brain tissue and LGG #p<0.05; normal brain tissue and HGG Δ p<0.05.

**Table 2 Different pathological grades of cerebral glioma MVD (**
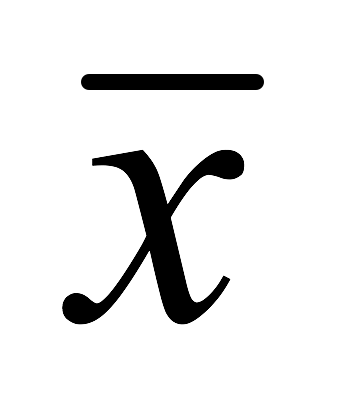
**±s)**

| **Grade** | **Cases** | **MVD** |
| --- | --- | --- |
| LGG | 26 | 47.4±5.7* |
| HGG | 23 | 63.5±7.6 |

Comparison between groups: LGG and HGG *p<0.05
